# Genome-wide identification of rhabdoviral sequences in alfalfa (*Medicago sativa* L.)

**DOI:** 10.64898/2026.05.20.726541

**Authors:** Sam Grinstead, Lev G. Nemchinov

## Abstract

We recently reported the identification of endogenous viral elements (EVEs) originating from the *Caulimoviridae* family within the alfalfa (*Medicago sativa* L.) genome. Our subsequent identification of ubiquitous rhabdoviral elements in infected and healthy alfalfa tissues by high throughput sequencing prompted us to suggest that the alfalfa genome might be populated with integrated rhabdoviruses as well. Bioinformatics analysis using 26 publicly available alfalfa genomes proved the suggestion accurate. We found multiple non-retroviral segments of the *Rhabdoviridae* family belonging to the genera *Betanucleorhabdovirus* and *Betacytorhabdovirus* that appeared to be stable constituents of the host genome. In that capacity they could potentially acquire functional roles in alfalfa’s development and response to environmental stresses. We believe this study reveals the first documented case of rhabdoviruses integrated into the alfalfa genome.

## Introduction

Endogenous viral elements (EVEs) are viral sequences integrated into host chromosomes and inherited by the host progeny as genetic alleles (Holmes, 2011; Bejarano et al. 1996; Nemchinov and Boutanaev, 2021). Plant genomes contain two main types of EVEs: endogenous pararetroviruses (EPRVs), which originate from reverse-transcribing dsDNA viruses of the family *Caulimoviridae*, and endogenous non-retroviral elements (ENREs) derived from ssDNA, ssRNA, and dsRNA viruses (Diop et al. 2018; Chiba et al. 2011; Takahashi et al. 2019; Chu et al. 2014). While integration of EPRVs into plant genomes is characterized relatively well (Staginnius and Richert-Pöggeler, 2006), insertion mechanisms of the ENREs remain largely obscure (Chiba et al. 2011). It is possible that pararetroviruses or retrotransposons provide the reverse transcriptase required to facilitate this integration (Chiba et al. 2011). As of today, no ENREs were documented in alfalfa.

We recently reported that alfalfa genome contains EVEs of the EPRV type and proposed that they are stable constituents of the alfalfa genome (Boutanaev and Nemchinov, 2021). Our subsequent discoveries of rhabdoviral sequences in both healthy and infected alfalfa plants via HTS indicated that the plant genome may contain integrated rhabdoviruses as well (Nemchinov et al. 2022, 2023ab, 2025ab).

In this work, using publicly available alfalfa genomes we performed bioinformatics analysis to identify the presence of endogenous viral elements, specifically those of the rhabdoviral origin. As a result of this study, many potential ENREs phylogenetically related to the genera *Betanucleorhabdovirus* and *Betacytorhabdovirus* were identified in alfalfa chromosomes 3, 4, and 5. As the findings were consistent across all utilized datasets/genomes, this may suggest that integrated rhabdoviral sequences carry a functional load similar to that of pararetroviruses, which have been linked to heritable RNA-mediated immunity in plants (Vali et al. 2023).

## Methods

A profile Hidden Markov Model (HMM) (Eddy, 1998, 2011) for the *Rhabdoviridae* family was constructed using HMMER v3.4 (http://hmmer.org/). Briefly, 85 complete plant *Rhabdoviridae* sequences were obtained from NCBI’s Refseq database, the nucleotide regions making up each conserved ORF (N, P, M, G, and L) (**Supplementary Table 1**) were aligned in MAFFT v7.055b (Katoh et al. 2002), and this alignment was used to generate a probabilistic model reflecting conserved family domains. The compiled HMM profile was then used to search for rhabdoviral-related EVEs in 26 alfalfa genomes (**Supplementary Table 1**; He et al. 2025, BioProject <u>PRJNA1220045</u>; and Zhang et al. 2025, BioProject <u>PRJNA1193221</u>). The E-value cutoff of < 1x10^-10^ was used to ensure low false-positive rates. Individual identified sequences were then searched via BLASTX against the ClusteredNR database to identify virus species. In a separate analysis, we searched the latest alfalfa genome annotation, ZM4_V2.0 (Zhang et al. 2025, BioProject PRJNA1179654), and extracted uncategorized regions marked as viral that lacked taxonomic classification. These sequences were then aligned against the NCBI non-redundant (nr) database using BLASTx to identify the specific virus species.

Phylogenetic analysis was performed with MEGA12.1 software (Kumar et al. 2024) using Maximum Likelihood method and Tamura-Nei model of nucleotide substitutions (Tamura and Nei, 1993).

## Results and Discussion

Screening 26 alfalfa genomes with Hidden Markov Model profiles revealed multiple rhabdoviral nucleocapsid protein (N) sequences across three chromosomes. Specifically, we identified sequences spanning 1,065–1,069 nt on chromosome 3 of five genomes, 1,016–1,022 nt on chromosome 4 of eight genomes, and 1,078–1,388 nt on chromosome 5 of all 26 genomes (**Table 1, Supplementary Table 2**). Individual analysis of all detected EVEs via BLASTX searches against the ClusteredNR database identified sequences that shared 78–100% identity (E-value=0; query coverage 89-100%) with the nucleocapsid proteins of two previously described viruses: alfalfa nucleorhabdovirus 1 (ANRV1, <u>UGO47147.1</u>) and alfalfa cytorhabdovirus 1 (ACRV1, <u>UGO47148.1</u>), (**Supplementary Table 3**), (Nemchinov et al. 2022). Phylogenetic analysis performed with MEGA12.1 software (16) clustered ANRV1 and ACRV1 EVEs found in 26 genomes with representative members of the genera *Betanucleorhabdovirus* and *Betacytorhabdovirus*, respectively (**Fig. 1**).

**Figure 1.**
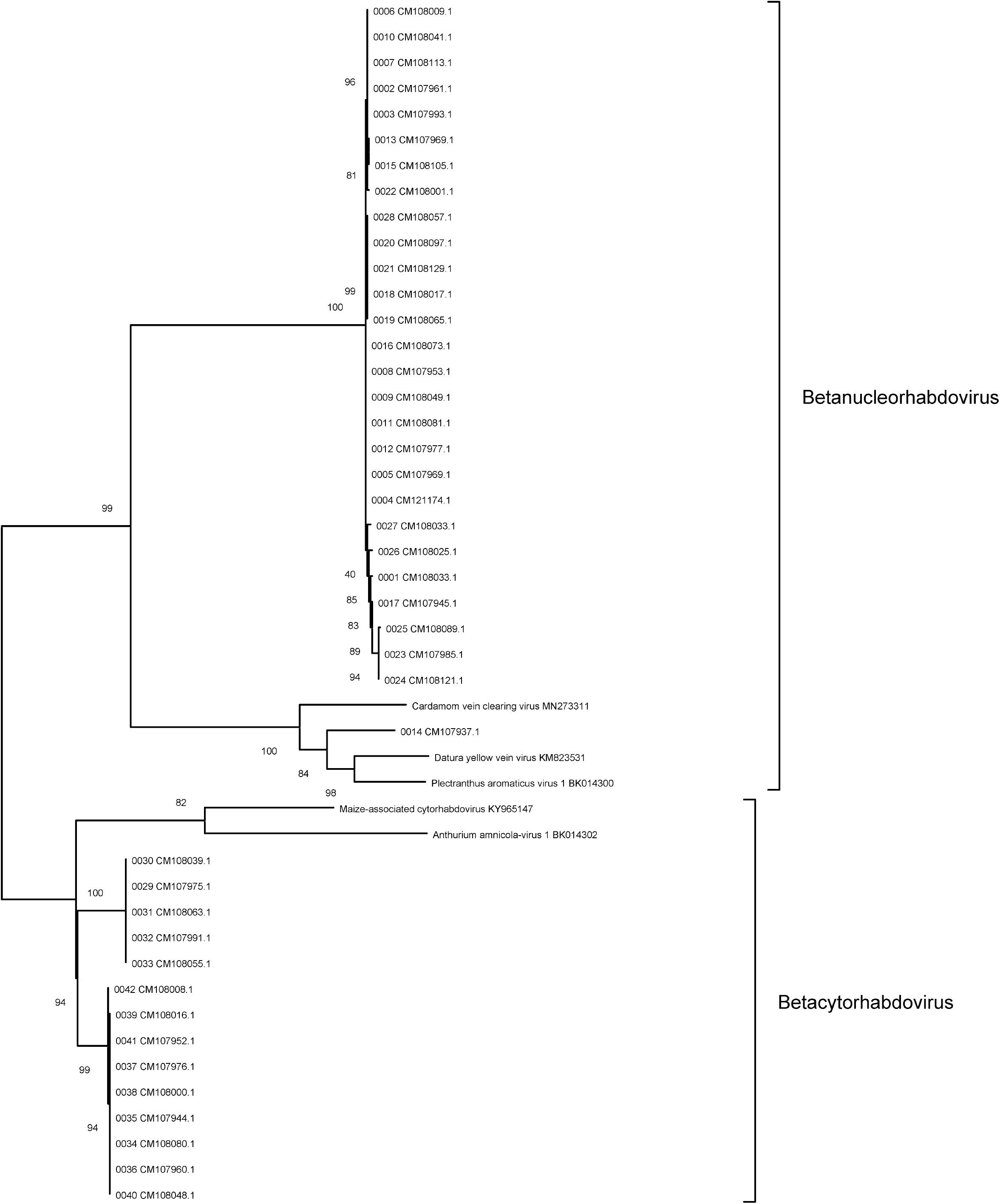
Phylogenetic tree depicting rhabdoviral sequences identified in 26 alfalfa genomes (<u>PRJNA1220045</u> and <u>PRJNA1193221</u>). The tree was constructed by MEGA 12 software (Kumar et al. 2024) using Maximum Likelihood method. The percentage of replicate trees in which the associated taxa clustered together (1,000 replicates) is shown next to the branches. The numbers 0001–0042 represent serial numbers, which are immediately followed by the corresponding accession numbers (e.g., CM108033.1).

**Table 1.** Rhabdoviral sequences identified in 26 alfalfa genomes (<u>PRJNA1220045</u> and <u>PRJNA1193221</u>).

| № | Accession | Query | Align from | Align to | E-value | Length | Target description |
| --- | --- | --- | --- | --- | --- | --- | --- |
| 1. | CM108033.1 | N* | 100591199 | 100589812 | 3.00E-118 | 1388 | M. sativa isolate S58 Chr 5 |
| 2. | CM107961.1 | N | 86335110 | 86333845 | 8.00E-117 | 1266 | M. sativa isolate S227 Chr 5 |
| 3. | CM107993.1 | N | 89087491 | 89086226 | 1.00E-116 | 1266 | M. sativa isolate S147 Chr 5 |
| 4. | CM121174.1 | N | 77730637 | 77729374 | 2.00E-116 | 1264 | M. sativa cv. Bolivia Chr 5 |
| 5. | CM107969.1 | N | 90220257 | 90218994 | 2.00E-116 | 1264 | M. sativa isolate S206 Chr 5 |
| 6. | CM108009.1 | N | 93133461 | 93132196 | 2.00E-116 | 1266 | M. sativa isolate S89 Chr 5 |
| 7. | CM108113.1 | N | 92244351 | 92243086 | 2.00E-116 | 1266 | M. sativa isolate B62 Chr 5 |
| 8. | CM107953.1 | N | 100444835 | 100443573 | 3.00E-116 | 1263 | M. sativa isolate S387 Chr 5 |
| 9. | CM108049.1 | N | 90584661 | 90583399 | 3.00E-116 | 1263 | M. sativa isolate S44 Chr 5 |
| 10. | CM108041.1 | N | 86457080 | 86455815 | 3.00E-116 | 1266 | M. sativa isolate S55 Chr 5 |
| 11. | CM108081.1 | N | 88733969 | 88732707 | 3.00E-116 | 1263 | M. sativa isolate B97 Chr 5 |
| 12. | CM107977.1 | N | 88724662 | 88723400 | 3.00E-116 | 1263 | M. sativa isolate S199 Chr 5 |
| 13. | CM107969.1 | N | 90898112 | 90896847 | 7.00E-116 | 1266 | M. sativa isolate S206 Chr 5 |
| 14. | CM107937.1 | N | 84162729 | 84163995 | 2.00E-115 | 1267 | M. sativa isolate ZM4 LG05 |
| 15. | CM108105.1 | N | 84414687 | 84413422 | 3.00E-115 | 1266 | M. sativa isolate B88 Chr 5 |
| 16. | CM108073.1 | N | 82032750 | 82031487 | 5.00E-115 | 1264 | M. sativa isolate B108 Chr 5 |
| 17. | CM107945.1 | N | 84122908 | 84121536 | 6.00E-115 | 1373 | M. sativa isolate ZM3 Chr 5 |
| 18. | CM108017.1 | N | 89226219 | 89224956 | 1.00E-113 | 1264 | M. sativa isolate S115 Chr 5 |
| 19. | CM108065.1 | N | 91424109 | 91422846 | 1.00E-113 | 1264 | M. sativa isolate B160 Chr 5 |
| 20. | CM108097.1 | N | 89006326 | 89005063 | 1.00E-113 | 1264 | M. sativa isolate B92 Chr 5 |
| 21. | CM108129.1 | N | 90152801 | 90151538 | 1.00E-113 | 1264 | M. sativa isolate B32 Chr 5 |
| 22. | CM108001.1 | N | 86771033 | 86769757 | 5.00E-112 | 1277 | M. sativa isolate S122 Chr 5 |
| 23. | CM107985.1 | N | 90950912 | 90949651 | 9.00E-112 | 1262 | M. sativa isolate S150 Chr 5 |
| 24. | CM108121.1 | N | 86108228 | 86106988 | 2.00E-111 | 1241 | M. sativa isolate B52 Chr 5 |
| 25. | CM108089.1 | N | 86792919 | 86791655 | 6.00E-111 | 1265 | M. sativa isolate B106 Chr 5 |
| 26. | CM108025.1 | N | 89328635 | 89327419 | 2.00E-107 | 1217 | M. sativa isolate S65 Chr 5 |
| 27. | CM108033.1 | N | 100592668 | 100591591 | 5.00E-105 | 1078 | M. sativa isolate S58 Chr 5 |
| 28. | CM108057.1 | N | 88346797 | 88345643 | 9.00E-101 | 1155 | M. sativa isolate S9 Chr 5 |
| 29. | CM107975.1 | N | 70112560 | 70113628 | 1.00E-56 | 1069 | M. sativa isolate S199 Chr 3 |
| 30. | CM108039.1 | N | 75075996 | 75077060 | 1.10E-56 | 1065 | M. sativa isolate S55 Chr 3 |
| 31. | CM108063.1 | N | 74001673 | 74002741 | 1.10E-56 | 1069 | M. sativa isolate B160 Chr 3 |
| 32. | CM107991.1 | N | 77960484 | 77961548 | 1.2E-56 | 1065 | M. sativa isolate S147 Chr 3 |
| 33. | CM108055.1 | N | 74364313 | 74365377 | 1.2E-56 | 1065 | M. sativa isolate S9 Chr 3 |
| 34. | CM108080.1 | N | 49500693 | 49501713 | 1.1E-54 | 1021 | M. sativa isolate B97 Chr 4 |
| 35. | CM107944.1 | N | 53841332 | 53842353 | 1.2E-54 | 1022 | M. sativa isolate ZM3 Chr 4 |
| 36. | CM107960.1 | N | 50846482 | 50847503 | 1.2E-54 | 1022 | M. sativa isolate S227 Chr 4 |
| 37. | CM107976.1 | N | 53913708 | 53914729 | 1.2E-54 | 1022 | M. sativa isolate S199 Chr 4 |
| 38. | CM108000.1 | N | 51479660 | 51480681 | 1.2E-54 | 1022 | M. sativa isolate S122 Chr 4 |
| 39. | CM108016.1 | N | 55348410 | 55349430 | 1.2E-54 | 1021 | M. sativa isolate S115 Chr 4 |
| 40. | CM108048.1 | N | 53332936 | 53333957 | 1.3E-54 | 1022 | M. sativa isolate S44 Chr 4 |
| 41. | CM107952.1 | N | 50836138 | 50837159 | 1.9E-54 | 1022 | M. sativa isolate S387 Chr 4 |
| 42. | CM108008.1 | N | 55864427 | 55865442 | 3.40E-54 | 1016 | M. sativa isolate S89 Chr 4 |

rhabdoviral nucleoprotein

When unclassified viral sequences from alfalfa genome ZM4_V2 were extracted (BioProject <u>PRJNA1179654</u>) and subjected to BLASTx search, several rhabdoviral EVEs were found in all four haplotypes of Chr 5 that mapped to ANRV1 (∼88–96% identity: E-value=0; **Supplementary Table 3**), partially confirming results of our 26-genome analysis (He et al. 2025).

As both ANRV1 and ACRV1 are rather phylogenetically distinct from other representatives of the family (Nemchinov et al. 2022), this may suggest that their EVEs are uniquely adapted to *Medicago sativa*. Consequently, they may have evolved and potentially acquire functional roles in alfalfa’s development and response to environmental stresses, as was reported for integrated pararetroviruses linked to heritable RNA-mediated immunity in plants (Vali et al. 2023).

We believe this study reveals the first documented case of rhabdoviruses integrated into the genomes of *M. sativa*. These findings also imply that some previously reported alfalfa rhabdoviruses may potentially represent EVEs rather than active infections, though verification is required.

## Author contribution

SG: concept, HMM-profiling, bioinformatics analysis. LGN: concept, phylogenetic analysis, first and last drafts of the manuscript. Both authors edited the final version of the manuscript.

## Funding

This study was supported by the United States Department of Agriculture, the Agricultural Research Service, CRIS number 8042-21500-003-000D.

## Ethics approval and consent to participate

No human participants were used in this study.

## Consent for publication

The authors consent to the publication of the manuscript.

## Conflict of interest statement

The authors declare no competing interests.

## Data availability

All sequences revealed and analyzed in this work are presented in Supplementary Tables.

## Notes

### Competing Interest Statement

The authors have declared no competing interest.

